# Plant height defined growth curves during vegetative development have the potential to predict end of season maize yield and assist with mid-season management decisions

**DOI:** 10.1101/2024.06.25.600633

**Authors:** Dorothy D. Sweet, Julian Cooper, Cory D. Hirsch, Candice N. Hirsch

## Abstract

Precision farming has been developing with the intention of identifying within field variability to adjust management strategies and maximize end of season yield and profitability and minimize negative environmental impacts. The development of quick, easy, and low cost methods to quantify field level variation is essential to successful implementation of precision agriculture at scale. Temporal plant height and growth rates collected with unoccupied aerial vehicles mounted with red, green, blue sensors have the potential to predict end of season grain yield, which could facilitate mid-season management decisions. Image-based plant height data was collected weekly from commercial maize fields in three growing seasons to assess variation within fields and the relationship with grain yield variation. Plant height, growth rate, and grain yield had variable relationships depending on the time point and growth environment. Models developed using temporal traits predicted grain yield variation within a commercial field up to r = 0.7, though insufficient water affected the prediction accuracy in one field due to the limited representation of drought environments in the model development. In the future, with more data from stress environments, such as drought, this method has potential for high accuracy grain yield prediction across a range of environmental conditions. This study demonstrates the potential of using unoccupied aerial vehicles to derive vegetative growth patterns and model within field variations, and has application in making mid-season management decisions.

## Introduction

Maize (*Zea Mays* L.) grain yields vary based on location, environmental conditions, and management practices. This variation occurs not only across and within regions, but also within individual fields. As a result, precision farming has developed in-part to quantify within-field spatial and temporal variability to adjust management practices and maximize end of season yield (Stafford, 2000). Adjusting management strategies based on sub-field resolution can benefit the environment by applying only necessary inputs, which will reduce the amount of run-off in water supplies. Adjustments can also benefit individual farmers and farming communities economics by preventing unnecessary costs in nutrient and pesticide application, and it can improve profitability and productivity by increasing grain yield and quality in areas of the field with higher intervention needs.

Early to mid-season estimates of yield could benefit multiple levels of the agricultural pipeline with farmers using the information to determine labor priorities and governments using the large-scale information for informed crop pricing, food trade policies, and food security (Hutchinson, 1991). Current methods of yield estimation before harvest vary widely in resolution, complexity, quantifiability, and need for crop expertise. The most common and traditional method is an estimation of yield based on expert farmer or agronomist knowledge of the field and the conditions present during that growing season through scouting efforts. This method results in a coarse estimate of yield that is flawed due to the inherently small sample size that could be collected through scouting, and requires substantial time, labor, and individual expertise. Use of destructive sampling on representative areas of the field can lead to slightly more accurate estimates, but the sampling process is both destructive and labor intensive. Loose estimates of the current yield based on the yield map of the previous year are also possible, but do not account for weather conditions in the current growing season.

Another possible approach to estimate grain yield variation takes advantage of the relationship between plant height and grain yield, as plant height has been shown to be predictive of maize grain yield in some regions and environments (Mallarino et al., 1999; Machado et al., 2002; Katsvairo et al., 2003; Yin et al., 2011; Barrero Farfan et al., 2013). Plant height is frequently used in research plots for assessing field plant vigor, estimating plant biomass, and predicting crop yield (Qiu et al., 2022). Studies focused on multiple crops have found that the relationship between plant height and grain yield can be augmented with machine learning to better predict yield. Machine learning with plant height data has been used to help predict yield in crops such as faba bean (Ji et al., 2022), winter wheat (Tao et al., 2020), and maize (Yang et al., 2021; Toledo et al., 2022) using different models and sometimes adding additional traits. Ji et al. (2022) compared support vector machine, random forest, and decision tree models and found that the yield estimation of support vector machine was the best, followed by random forest, and then decision tree. In winter wheat, Tao et al. (2020) used partial least squares regression, random forest, and an artificial neural network to predict yield using spectral indices, ground plant height, and hyperspectral measured plant height and found the partial least squares regression model, when trained using a combination of the spectral indices and hyperspectral extracted plant height, had the best predictions followed by artificial neural network then random forest.

Manual measurements of plant height using a ruler are time consuming and labor intensive, which is why much research has been put toward extracting plant height from images collected by unoccupied aerial vehicles (UAVs), especially with red, green, blue (RGB) sensors due to their simplicity, availability, and affordability (Araus and Cairns, 2014; Yang et al., 2017; Madec et al., 2017; Tirado et al., 2020). While there is great potential in applying UAVs to measure plant height in commercial production fields to predict end of season performance, this approach has been challenging to implement in commercial fields due to the lack of plot structure used in ground subtraction methods (Fujiwara et al., 2022), the necessity for ground control point (GCP) placement for accurate structure from motion plant height extraction (Rosnell and Honkavaara, 2012; Wu et al., 2020; Fujiwara et al., 2022), and limitations on the distances UAVs can fly due to federal regulations and battery life (Sweet et al., 2022).

Other sensors and platforms have been used with varying degrees of success to measure plant height in production fields. In a commercial soybean field, tractor-mounted ultrasonic distance sensors were used to measure plant height on four rows of soybeans simultaneously and obtained a height accuracy within 2 cm (Fletcher and Fisher, 2019). While this method was effective in a soybean field, collecting measurements 4 rows at a time is time-consuming and as maize gets much taller throughout the seasons measurements would be increasingly difficult to use this approach. In commercial cotton fields, plant height was estimated using satellite imagery containing multi-spectral bands and machine learning, with height estimates over large areas, averaging 15 measurements across 175 acre fields (da Silva Andrea et al., 2023). While this study was able to show some success in estimating plant height with R^2^ values as high as 0.87, the study areas are too large for useful sub-field resolution information needed for precision farming. Similarly, a study in maize using satellite imagery from Sentinel-1 to estimate plant height found that, although the average height across the field was the same as the average of the manual measurements collected on the ground, there was large across field variation that did not reflect real values, with errors of around 50 cm (Arslan et al., 2022). This method had a large error rate, and even when accurate had insufficient spatial resolution to be useful for determining within field variation. As these studies have shown, tractor-mounted sensors are too slow and low to the ground to effectively capture maize growth rates throughout the season; while affordable satellites have a spatial and temporal resolution that is too large to effectively show sub-field plant height variation. In contrast to these platforms, development of UAV mounted RGB sensors for production fields would offer an inexpensive platform that can be flown at very high temporal resolution. UAVs are flown above the canopy which allows their use with taller crops like maize, while also not being too high to allow for high within field spatial resolution.

Calculating plant height values using UAV images has been achieved using multiple methods for ground estimation including digital terrain model interpolation, exposed alley subtraction, manual measurement self-calibration, and a difference based method (Sweet et al., 2022). These methods were originally developed for use in research fields with plots and alleys and most of them require defined plots or visible soil in between plants, which will limit their use for commercial fields, especially later in the season when the canopy is mostly closed. For instance, the digital terrain model interpolation method creates a digital terrain model by identifying bare-ground pixels and masking out plant pixels. The masked values are filled in with estimated ground values. Finally, the digital terrain model is subtracted from the digital surface model (DSM) with the plant values included to obtain plant canopy height (Bendig et al., 2014; Holman et al., 2016; Watanabe et al., 2017; Varela et al., 2017; Chang et al., 2017; Madec et al., 2017; Malambo et al., 2018; Thompson et al., 2019; Hassan et al., 2019; Han et al., 2019). This method does not require plots, but necessitates visible ground which may not be available in a commercial maize field, especially later in the season.

In contrast, the difference based method does not require plots or visible ground to calculate plant canopy height. In this approach a DSM from a flight completed before the crop is visible is used as the ground value while a DSM from a flight on the day of interest is used as the plant canopy values. The ground values are then subtracted from the plant canopy values to obtain plant height measurements (Chu et al., 2018; Acorsi et al., 2019; Belton et al., 2019). This method relies on accurate flight paths, DSM registration, high DSM resolution, and smaller pixel sizes for improved accuracy, which can increase computational load. Despite these limitations, the difference based method provides a practical solution for ground estimation when using UAV collected RGB images to calculate sub-field plant height variation in a commercial maize field.

To assess the utility of UAV mounted RGB sensors for precision maize farming, we (1) obtained RGB images of maize production fields across three seasons using a UAV mounted RGB sensor, (2) developed methodology to accurately extract plant height from these images throughout vegetative development, (3) evaluated the relationship between plant height and yield in variable environmental conditions, and (4) assessed the ability to predict grain yield using machine learning models trained on plant height, plant height estimated growth rate, and RGB based vegetative indices. We demonstrate the ability to accurately extract high resolution plant height data from commercial production fields, and the potential to use this data for modeling end of season grain yield, facilitating precision farming efforts.

## Materials and Methods

### Field Setup

To ensure the field conditions replicated true commercial field practices, existing maize commercial production fields planted for grain harvest and subject to typical management practices were used. The fields were located in Waseca, MN 44.0770° N, 93.5084° W in 2020-2022. Each year the fields were planted with a current commercial hybrid (2020: Channel 199-11 STXRIB; 2021: Becks 5077 V2P; and 2022: LG 5525 VT2 Pro) at 35,500 seeds/acre; and fertilizer was applied as specified in Supplemental Table S1. The entire planted field was large and only a subset of the field (∼ 3 acres) was evaluated to allow for sufficient manual measurements distributed throughout the flight area to be collected in the same day.

### UAV Data Collection and Processing

An initial flight of the field each year was collected before emergence to capture ground topography for use in ground subtraction calculations. After emergence, flights were conducted approximately every week until plants reached terminal height, defined by anthesis, using a DJI Phantom 4 RTK drone. Images were collected at an altitude of 30 m above the ground. This achieved a ground sampling distance of approximately 0.82 cm with 80% front overlap and 80% side overlap early in the season and 85% front and side overlap later in the season to maximize reconstruction efficiency. A UAV base station was placed at the same location on the outside border of the field for each flight within the year and the GPS location was set to the same coordinates for each flight. Flights to measure plant height (not counting the initial ground flight) were collected at 7 timepoints in 2020, 6 timepoints in 2021, and 7 timepoints in 2022 (Supplemental Table S1). The ground targets were two feet in width and were placed around the border of the defined ‘field area’, meaning that some were placed in the middle of the larger production field to be used as GCPs for the flight area (Supplemental Figure S1). PVC pipes were used as posts for the GCPs and the height of the posts varied as the season progressed to stay above the canopy. The posts were 5 ft long at the beginning of the season and by the end of the season they were extended to 10 ft. The real world coordinates of the GCPs were collected using real time kinematic positioning with a Swift Console (v 2.3.17) base station and rover (GNSS compass configuration,). As the height of the GCPs changed, height (m) was added to the altitude measurement of the GPS location (Supplemental Table S2).

### Manual Measurement Data Collection

Randomly distributed plots that were 10 feet long and one row wide were defined throughout the field to collect manual height measurements at consistent locations throughout the growing season (40 in 2020, 20 in 2021, and 20 in 2022) (Supplemental Table S3). Manual plant height measurements were collected within these defined plot areas on the same day as each drone flight (Supplemental Table S1). Within a plot, hand measurements were collected on five representative plants from the center of the plot using a ruler. During vegetative growth, plant height was measured as the distance between the ground and top most freestanding vegetative part of the plant. At reproductive maturity plant height was measured as the distance between the ground and the top of the tassel. The manual measurements were used to correlate with extracted height values from the same plot to determine the relative accuracy of the extracted height values.

### Image Processing

Images from each flight were processed as previously described (Tirado et al., 2020) until reaching the height extraction step, at which point a different MATLAB script was used (Supplemental Figure S2) (Inc., 2022). Briefly, Agisoft Software (Agisoft Metashpe Professional v1.7.5) was used to process the images to generate crop surface models and RGB orthomosaics for each flight, including the initial ground flight. The QGIS software (QGIS v3.16, 2021) was used for plot boundary extraction by overlaying a grid of squares based on the width of 6 rows (120 inches) and exporting plot coordinates (Supplemental Figure S1). A width and length of 120 inches was used for the grid size as these are dimensions of realistic management intervention. Plot boundaries of 120 inches long and 29 inches wide were also extracted for the areas defined as manual measurement plots in the field (Supplemental Figure S1).

### Plant Height Extraction, Normalization, and Quality Control Analysis

The difference-based method (Chu et al., 2018; Acorsi et al., 2019; Belton et al., 2019) was applied to extract plant heights for the plot squares using MATLAB scripts that are available in the GitHub repository linked in the Data and Code Availability section below (Supplemental Figure S2) (Chu et al., 2018; Acorsi et al., 2019; Belton et al., 2019). Using this method, height was extracted for each 10ft-by-10ft square of the grid (Supplemental Figure S1) from a DSM of the initial ground flight (extracting the 3rd percentile of the height value from all the pixels in a given square of the grid) and from a DSM of the data flight (extracting the 97th percentile of the height value from all the pixels in a given square of the the grid). The height of each plot boundary at the time of the data flight was determined by subtracting the initial ground flight height value from the data flight height value. This method was similarly used to extract height values for the manual plots and plots encapsulating the GCPs (Supplemental Figure S1). All of the raw extracted plant height values are stored in the DRUM repository linked in the Data and Code Availability Section.

The extracted plant heights, both the whole field grid plots and the defined manual plots, were normalized to real world measurements by comparing the extracted GCP heights to the known GCP heights (Supplemental Figure S3). This was completed using three different methods depending on the specifics of each flight, and if (i) the height of the UAV base station was able to be extracted from the DSM (Supplemental Table S4), which was not always the case due to the small size of the base station, and (ii) the ground measurement on a given flight differed from the pre-emergence ground flight. The first two normalization methods corrected for differences in ground measurements between the initial ground flight and the data flight. The first method extracted a height value from the bare ground to determine an overall zero value for the flight and extracted a height value from the UAV base station to establish a precisely known upper height. The second method also extracted a height value from the bare ground to determine an overall zero value for the flight, but extracted a height value from the GCPs to establish a precisely known upper height when a height was unable to be extracted from the UAV base station. The third method was applied when a ground correction was not required, and only a height value for the UAV base station was extracted for a precisely known upper height. No matter the extraction method, height normalizations were completed by adjusting all the heights to the ground value with subtraction (when it was necessary). Then the values were further adjusted to real world heights by obtaining the ratio of real world upper height to extracted upper height for either the UAV base station or GCPs, based on the particular flight, to get the adjustment factor. This factor was used to multiply with the extracted plant height values for each plot to obtain normalized plant heights for each plot. Pearson correlations between the normalized extracted values and manual height measurements were completed to validate the extracted values (Supplemental Figure S4).

Individual data points (i.e. single grid squares within a single flight date) were removed from the dataset if they were classified as a dip, defined as having a height less than 80% of the previous flight day and also remained less than the next flight day, or a peak, defined as having a height more than 120% of the next flight day while still remaining more than the previous flight day flight. In total, 618 data points (4 for 2020, 70 for 2021, and 544 for 2022) out of 24,129 total data points (9,380 for 2020, 6,720 for 2021, and 8,029 for 2022) were removed based on these criteria (Supplemental Figure S3, Supplemental Table S5).

### Color Value Extraction and Vegetative Index Calculation

To calculate vegetative indices the plot pixels were classified based on raw RGB values using a k-means classification algorithm implemented in MATLAB (Inc., 2022). Clusters were manually identified as containing plant material or background noise such as soil and shadow. Clusters containing background noise were removed from further analysis. The clusters that contained plant pixels were masked within each orthomosaic and dilated using the ‘imclose’ function in MATLAB to fill in small areas of plant pixels within plant clusters that were grouped into non-plant clusters. Red, green, and blue color values were extracted from the masked RGB orthomosaics on a plot basis using the same plot overlay as used for plant height. The color values for all the masked pixels in a plot were averaged to create a single value for each color channel. From these values, 13 vegetative indices were calculated (Supplemental Table S6).

### Yield Data Collection and Calculation

Yield was collected using a yield monitoring system attached to a combine with a 6-row head that was used for harvesting. Each yield data point was associated with GPS coordinates and exported into .shp and .csv files. These data were then associated with the plant height extraction plot squares by locating the 9 yield data points closest to the center of each plot square and calculating a weighted mean based on the distance of each data point from the center of the plot square. In 2021 and 2022, a line of yield data was removed from the right side of the field where the combine turned around each pass after removing the border rows. These data were removed due to artificially low yields resulting from the combine head running for a short time before any grain was harvested which gave the impression of more area having been covered than was actually covered.

### Weather Data and Growing Degree Day Calculations

Weather information including daily minimum and maximum temperatures (°F), total precipitation (in.), and total solar radiation (Cal/cm^2^) were collected from the University of Minnesota Southern Research and Outreach Center weather station (Station ID: 218692) in Waseca, MN, which is located less than 1 mile from the field locations. Growing degree days (GDD) were calculated for each day using the equation: *GDD* = [(*Tmax* + *Tmin*)/*2*] − *50*, where GDD is the growing degree days accumulated for an individual day and Tmax and Tmin and the maximum and minimum recorded air temperatures for the day. Temperatures above 86°F were adjusted to 86°F and temperatures below 50°F were adjusted to 50°F as corn growth rates do not increase or decrease outside of this range. Individual flight dates were converted to the cumulative sum of GDD from the planting date to the flight date.

### Curve Fitting using LOESS Regression Modeling

Separate LOESS regression models were fit to the cleaned, normalized plant heights for each plot square using all remaining flight time points using the ‘loess’ function in the ‘stats’ package v.4.2.2 in R v.4.2.2 (R Core Team, 2019). A span was chosen for the regression model for each year which varied between 0.4 - 0.6 based on an error matrix developed with 100 folds to determine the best fit (Supplemental Figure S5). Plant height values for every 50 GDD of the growing season for each grid square within a year were predicted using the fitted model. On average, 50 GDD equated to approximately 2.8 calendar days. The rate of growth between two time points was calculated as the slope between the two values.

### Correlation Analysis

The Pearson correlation between yield and other measured and derived traits was calculated using the ‘cor’ function in the ‘stats’ package v.4.2.2 in R v.4.2.2 (R Core Team, 2019). Correlations were calculated for each year individually, as well as for all three years together.

### Regression Analysis and Partial R^2^ from Multivariate Regression Models

Linear regression models with yield as the response variable were created for testing using the ‘lm’ function from the ‘stats’ package v.4.2.2 in R v.4.2.2 (R Core Team, 2019). A separate model was created for each data type (i.e. plant height and growth rate) and each year, with all terms treated as fixed effects which resulted in six separate global models (b ∼ t_1_ + t_2_ + t_…_+t_n_ + ε where b is grain yield in bu/acre, t is the phenotypic trait (plant height or growth rate) for each time point (1 to n), n is the number of time points, and ε is residual). Partial R^2^ for each regression model was calculated using the ‘partial_r2’ function from the ‘sensemakr’ package v.0.1.4 in R v.4.2.2 (R Core Team, 2019; Cinelli and Hazlett, 2020).

Each global model was passed through the ‘dredge’ function from the ‘MuMIn’ package v.1.47.1 in R v.4.2.2 (R Core Team, 2019; Bartoń, 2023) to determine the explanatory variable combination with the lowest second-order Akaike information criterion (AIC_c_). AIC_c_ was calculated using the following formula:

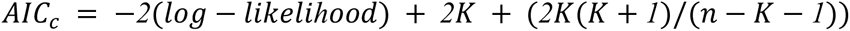

where n is the sample size and K is the number of model parameters. The data was split by data type for ease of comparisons and to restrict the number of explanatory variables required due to computational limitations. The relative importance value for each predictor remaining in the combination with the lowest AIC_c_ was calculated using the sum of weights function ‘sw’ from the ‘MuMIn’ package v.1.47.1 in R v.4.2.2 on the output of the ‘dredge’ function (R Core Team, 2019; Bartoń, 2023).

### Functional Principal Component Analysis

Functional principal component analysis (FPCA) was used to reduce the dimensionality of the highly correlated data extracted at 50 GDD windows. The FPCA for the curves was calculated using the ‘fda’ package v.6.0.5 in R v.4.2.2 (Ramsay et al., 2009; R Core Team, 2019; Rickert, 2021). The LOESS curve data was first put into a format able to be understood by the ‘pca.fd’ function using the ‘Data2fd’ function. Then, an object of class pca.fd was obtained from the function ‘pca.fd’ that contained the eigenvalues (values), the eigenvectors (harmonics), the proportion of variation explained by the eigenvalues (varprop), a value to rate how well each plot fit each eigenvector (scores), and other parameters. The top two eigenvectors were used as they explained more than 90% of the variation found in the LOESS curves. The eigenvalue for the first two eigenvectors was kept as data labeled functional principal component 1 (FPC 1) and 2 (FPC 2).

FPCA was used to represent vegetative indices throughout each season to compare them across years, as flights were not collected on the same day each year and curves could not be fit to these non-continuous values. The same method as described above was used, which resulted in two FPC scores for each plot in each year. The number of plots was 1340, 1120, and 1147 in 2020, 2021, and 2022 respectively, and combined with each of the calculated 16 vegetative indices resulted in a total of 115,424 data points of which 107,392 remained after quality control removal of plots.

### Machine Learning

Machine learning algorithms were used to predict grain yield using feature combinations of temporal plant height, temporal growth rate, the slope of the exponential growth, the GDD of the start and end of the exponential growth, the top two FPCA scores to describe the full growth curve, and the top two FPCA scores to describe each RGB vegetative indice for each plot (Table 1). The generation and evaluation of models was done using the ‘caret’ package (Kuhn, 2008) v.6.0 - 93 was used in R v.4.2.2 (R Core Team, 2019). The initial models trained on all of the data features included lasso, pcaNNet, pls, plsRglm, rf, svmRadial, and svmPoly. These were reduced down to pls, plsRglm, and rf based on initial model performance. For all feature combinations, the data was split into training and validation sets based on year, resulting in 3 different training and validation pairings (2020 and 2021 training for 2022 validation, 2020 and 2022 training for 2021 validation, and 2021 and 2022 training for 2020 validation) using an approximate 66/33 percent split for each combination. This method of separating the data was used due to the desire to predict yield in new environments (both geographical and weather) with yield data from other environments. The different models were trained with a list of hyperparameters using the training argument tuneGrid (1:21 for ncomp in pls; 1:20 for nt and (0.001, 0.01, 0.1, 1) for alpha.pvals.expli in plsRglm, and 2:11 for mtry in rf. To increase the generalization of the model, cross validation was completed with fold assignment based on year.

**Table 1.** Phenotypic features included in machine model models. All features included in each iteration of model testing and the associated title assigned to that combination of features. Numbers in parentheses indicate the number of features of each type.

| Data Combination Title | Feature Components |
| --- | --- |
| All Data | Temporal plant height (16), temporal growth rate (15), slope of exponential growth (1), GDD of start of exponential growth (1), GDD of end of exponential growth (1), top two FPCA scores for full growth curve (2), and top two FPCA scores for each RGB vegetative indices (32) - total: 68 features |
| Height, Rate, Eigen | Temporal plant height (16), temporal growth rate (15), slope of exponential growth (1), GDD of start of exponential growth (1), GDD of end of exponential growth (1), top two FPCA scores for full growth curve (2) - total: 36 features |
| Height, Rate | Temporal plant height (16), temporal growth rate (15) - total: 31 features |
| Growth Eigen | Top two FPCA scores for full growth curve (2) - total: 2 features |
| Vegetative Index Eigen | Top two FPCA scores for each RGB vegetative indices (32) - total 32 features |

The scaled variable importance was calculated for each model - data type combination using the ‘varImp’ function with scale set to true to keep the analysis comparable across model types.

### Data and Code Availability

All of the scripts and files used to generate and analyze data are available on GitHub at https://github.com/HirschLabUMN/Production_Drone_Height.git. All of the data that includes the UAV-derived plot height values, UAV-derived vegetative index values, orthomosaics, DEMs, plot boundary files, and mask files for each date of UAV data collection, cumulative GDDs calculated for each date of data collection, manual height data, combine harvested grain yield data, field management information, soil data, and weather data is available at the Digital Repository for U of M (DRUM) at https://doi.org/10.13020/7t39-h236.

## Results and Discussion

### Plant height can be accurately extracted from RGB images in commercial maize fields

To determine if accurate plant height estimates could be obtained from an UAV equipped with a RGB sensor, images were acquired from commercial production fields over three growing seasons (Figure 1A-C). The difference based method was used to estimate average plant height in 10ft-by-10ft squares throughout the growing season (Supplemental Figure S1). The distribution and shape of the growth curves of the extracted plant heights across the growing season varied from year to year (Figure 1D). This difference both within and between fields is more readily seen in calculated growth rates than the trait data per se. To assess the accuracy of the extracted height measurements, manual measurements in a subset of the plots were obtained (Supplemental Figure S4), and the Pearson coefficient between the normalized extracted plant height values and the manual plant height measurements were calculated. The correlation coefficient between normalized extracted height and manual heights across all flights within a field and year ranged from an R^2^ of 0.84 to 0.94 (Figure 1E-G). In other studies conducted in research fields with defined plots and alleys the correlation of image extracted heights with manual measured heights across flights within a year achieved comparable correlations (Varela et al., 2017; Madec et al., 2017; Tirado et al., 2020). The correlations across the growing season contained data from multiple growth stages with a large variance between them, which can inflate the overall correlation. Within specific flight dates the correlations were lower overall (R^2^ of 0.01 to 0.77; Supplemental Figure S4). Early flights for each year and location showed the lowest correlation between extracted and manual heights due to the low variation across the field at this early stage, which mathematically limits larger correlation values. This is similar to results from other studies with lower correlations achieved on individual flight dates, especially early in the season (Tirado et al., 2020; Volpato et al., 2021). Late season flights also had challenges due to field reconstruction issues. Flights later in the season required there to be nearly no wind for accurate image stitching, particularly when there were no outside edges and minimal identifiable features in the field to assist the stitching software. In 2022, all late flights were collected on very still days, and able to be stitched through flowering, but this was not the case for 2020 and 2021 flights.

**Figure 1.**
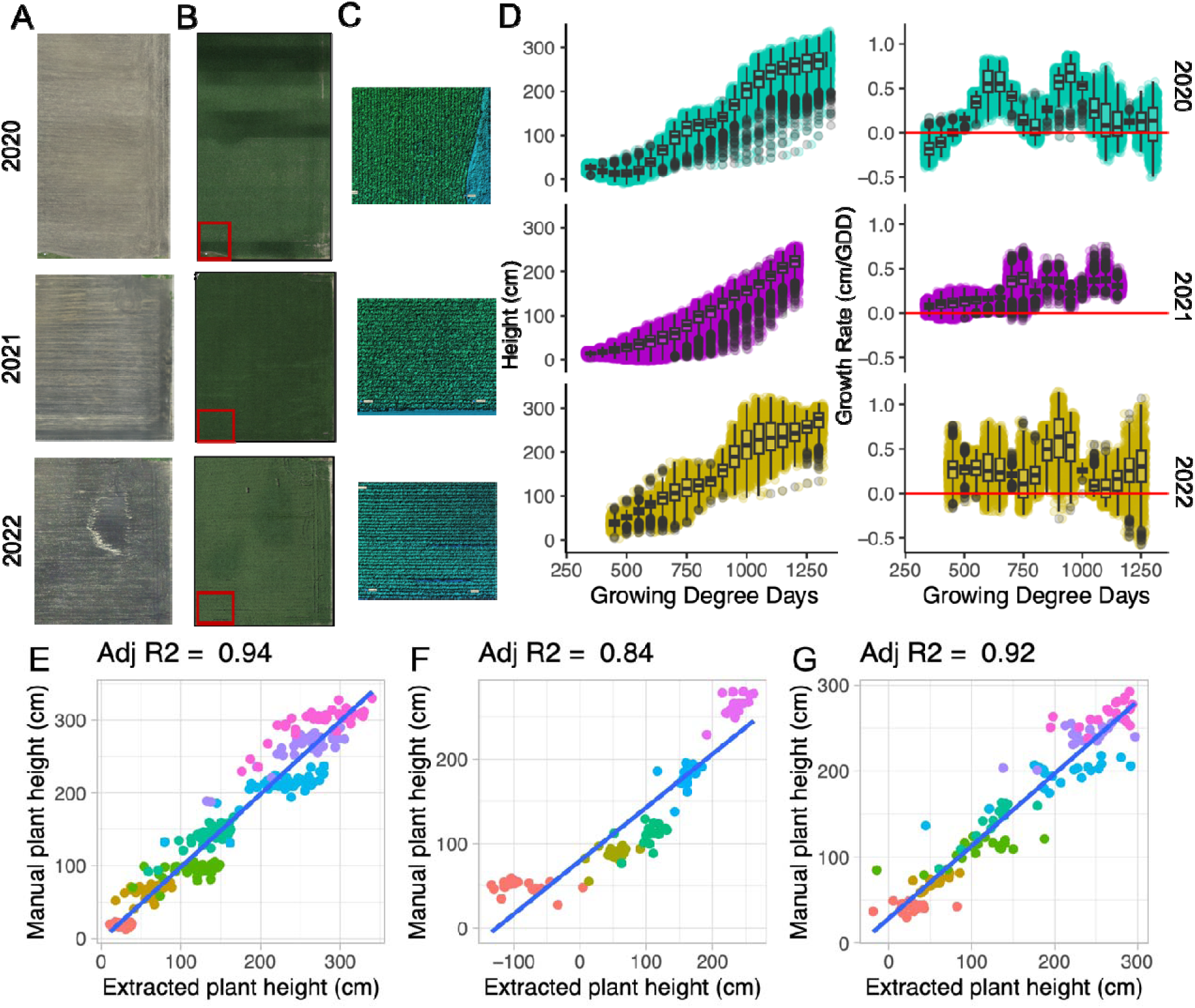
Plant height and growth rate extraction. (A) Ground flight orthomosaic for each year (05212020, 05182021, 05262022). (B) Example data flight orthomosaic for each year (07202020, 07122021, 07182022). The red box indicates the portion of the field shown in part C. (C) Example of Digital Elevation Model (DEM) showing height variation within the field for each year. (D) Extracted plant height fit to loess curves over the growing season and growth rates across time for each year. Pearson correlation between mean extracted plot plant height and mean manual plot plant height measurements across all dates within (E) 2020, (F) 2021, and (G) 2022 for height extraction validation. Different colors in E-G indicate data from different flight dates within each year.

### Plant height and growth rate have variable correlation with yield across the growing season and across growth environments

To assess the relationship of these extracted plant height and growth rates with yield, a yield map was obtained from the harvest combine (Figure 2A-C). In 2020, the yield across the top and right sides of the field, as well as a large elongated spot on the left side were lower than the rest of the field (Figure 2A). In 2021, there was lower yield in the bottom left portion of th field as well as the top left and the corners of the bottom and top right of the field (Figure 2B). The yield in 2022 was fairly consistent with very little variation across the portions of the field that were able to be kept after plant height quality control (Figure 2C). The 2021 and 2022 data were collected from the same field, yet substantial differences in the spatial patterning of yield were observed. In all cases, the yield distributions were consistent with visual assessments of the plants in the field throughout the season and prior seasons yields. The yield map was used to estimate the grain yield of each of the extracted plot squares based on a weighted mean that similarly reflected the differences in performance within and between years (Figure 2D). This variability in performance across years was desirable for this study to determine if the models were generalizable with predictions that are able to extend across different growth environments.

**Figure 2.**
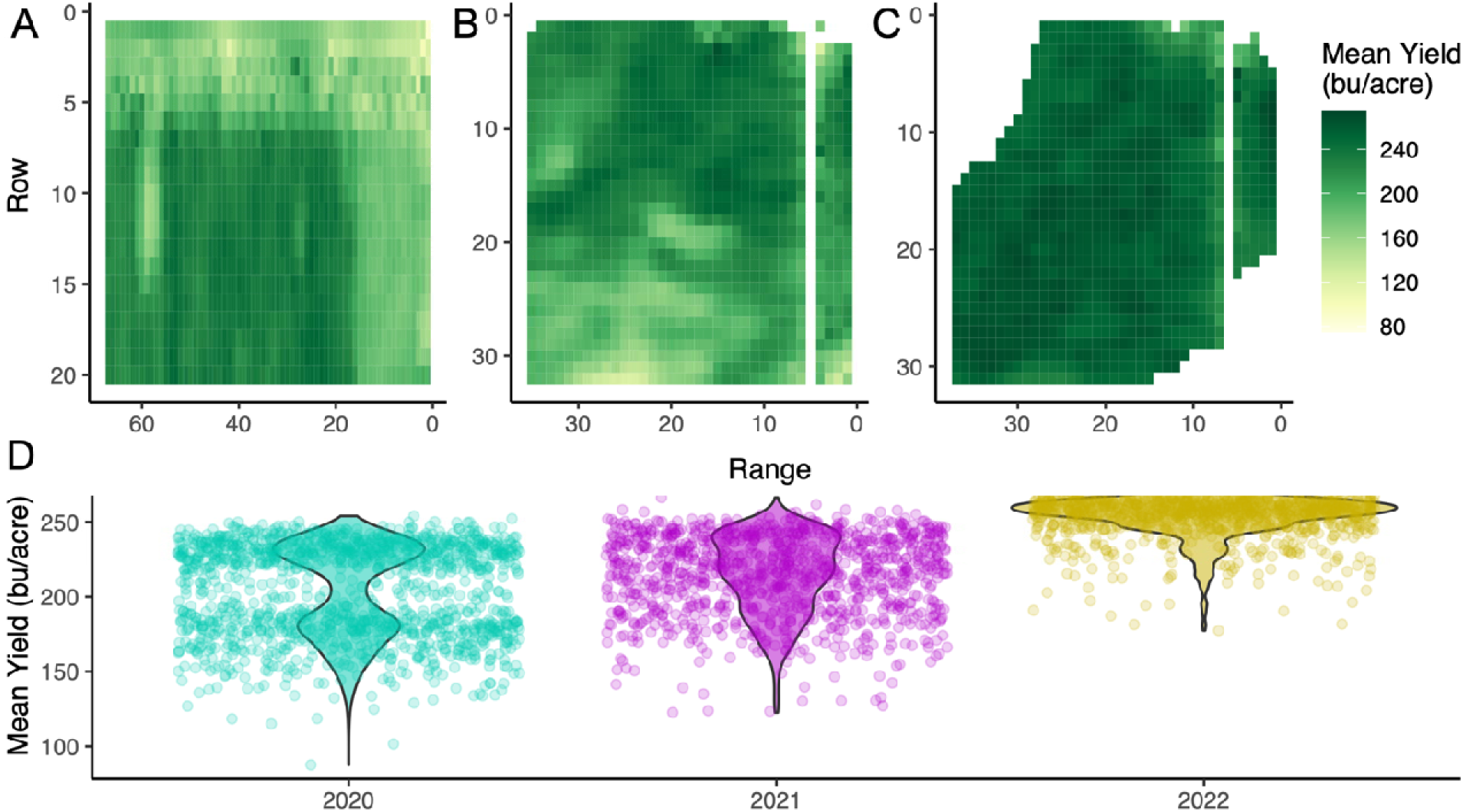
Grain yield variation. Heatmap of grain yield variation within the commercial maize field in (A) 2020, (B) 2021, (C) 2022. (D) Distribution of mean grain yield across extracted plots for each year. In 2021 and 2022, a line of yield data was removed from the right side where the combine turned around each pass after removing the border rows.

Pearson correlation coefficients between plant height and growth rate at each GDD and final yield showed very different results across the 3 environments (Figure 3A). Throughout 2021 there was little to no correlation between any of the plant height or growth rate time points and yield with the strongest relationship of r = −0.36 at 1150 GDD for growth rate. In contrast, stronger consistent correlations were observed across 2020 and 2022 (r = 0.76 at 1000 GDD for growth rate in 2020 and r = 0.61 for 850, 950, and 1050 GDD for growth rate in 2022). The growth rates at 900 to 950 GDD consistently had a positive correlation with yield across all three years. The correlations in 2020 (r = 0.7) and 2022 (r = 0.6-0.61) were more robust, but 2021 still had its strongest positive correlation during this time point (r = 0.29-0.35). This time point is during the early exponential growth phase, where a positive correlation with yield makes sense as faster growing plants during this time frame are likely to have an advantage over slower growing plants and often result in a taller overall height. In contrast, the 1050-1200 growth rate has a consistently negative correlation with yield, although not as robust as the positive correlation earlier in the season. A negative correlation with growth rate makes sense at this later point in the season as plants do not continue to grow after reaching the vegetative to reproductive transition (Abendroth, 2011). If any plants are still growing quickly in this stage they are likely behind in development and unlikely to produce as much. Just as there was an expected negative correlation between growth rate and yield late in the season, we would expect a stronger positive correlation between plant height and yield towards the end of the growing season as taller plants within a genotype usually have a higher yield (Wang et al., 2018). Between 900 and 1100 GDD, plant height had a positive correlation with yield in both 2020 (r = 0.65-0.74) and 2022 (r = 0.46- 0.58) that increased toward the end of the season.

**Figure 3.**
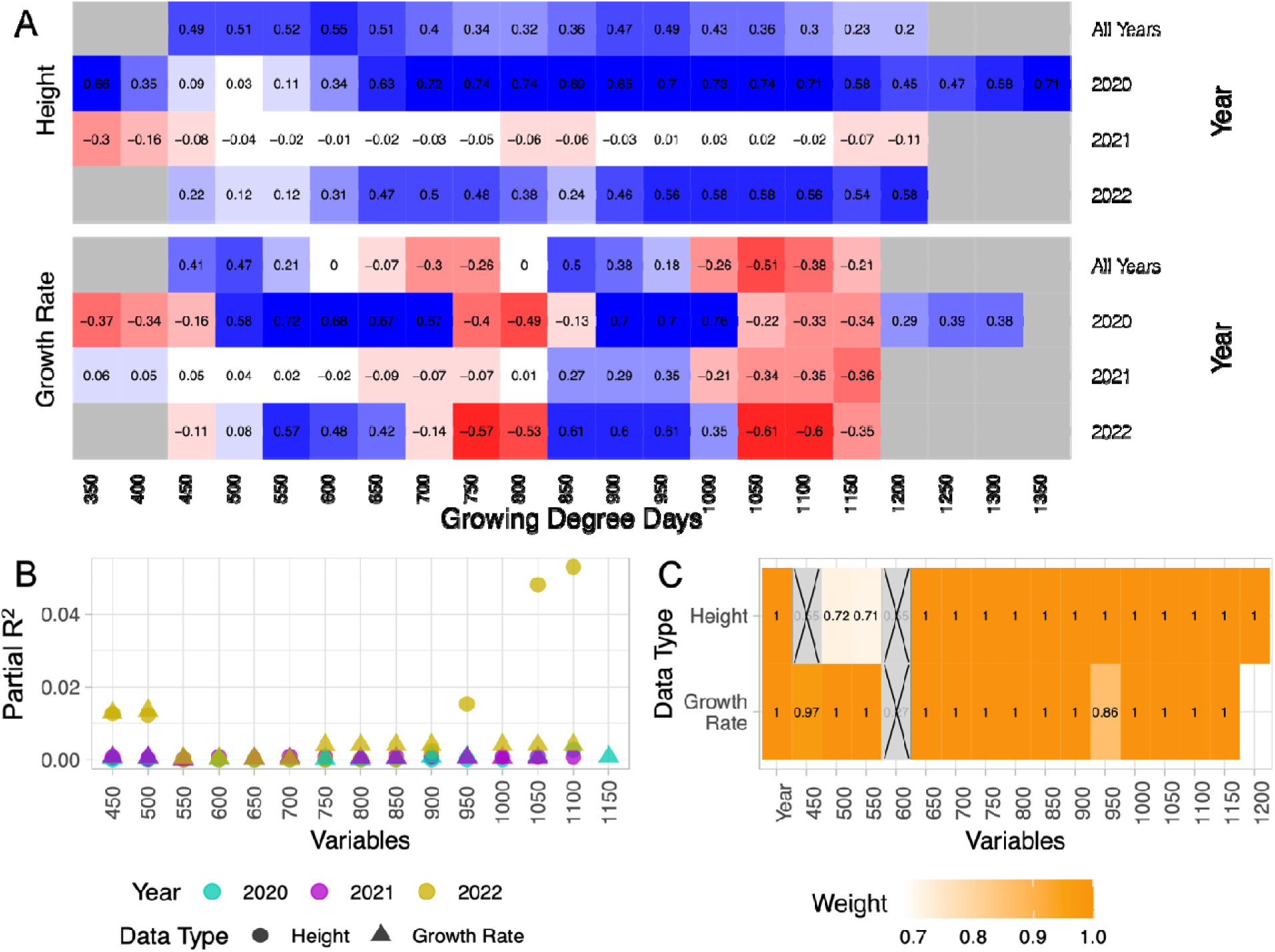
Relationships and relative importance of temporal traits. (A) Pearson correlation coefficient (r) between grain yield and plant height (top) and growth rate (bottom) for individual years and all years together. Gray boxes indicate missing data. (B) Partial R^2^ values for temporal variables in plant height and growth rate for each year. (C) Relative importance and retention of variables in AIC selection regression models.

To further understand the relationship between plant growth throughout development and end of season yield, partial R^2^ was calculated using a model that included plant height or growth rate for each GDD. A higher partial R^2^ indicates that an explanatory variable (i.e. a specific time point in development) explains more of the variance in the response variable (i.e. end of season yield). Partial R^2^ for plant height and growth rate variables across the season were very similar except at 1050 and 1100 GDD in 2022, when plant height had a substantially larger partial R^2^ (R^2^ = 0.004 vs. 0.05) (Figure 3B). No single time point throughout the growing season had a large partial R^2^ value, although some variables, such as 550 and 700 GDD, had near zero partial R^2^ values for all data types and years.

To better understand the importance of each explanatory variable, and the efficiency of including each in a final model, an AIC selection regression model was used. A lower AIC_c_ value for a model indicates a better chance of predictive success with a separate dataset and a higher weight within the model indicates a larger contribution of the explanatory variable. The weights of each explanatory variable in the AIC selection regression models aligned closely with the results of the partial R^2^ values. The variables with the lowest partial R^2^ values also had the lowest relative importance and some were even removed from the final model (Figure 3C). The lowest partial R^2^ values for plant height were from 450 to 700 GDD in 2020 and 2021 and from 550 to 700 in 2022. Plant height at 450 and 600 GDD were removed during model selection, while plant height at 500 and 550 were retained, but had lower weights in the final model than other variables. The first 4 models all had a delta AIC lower than 2, indicating other model combinations with variation on the inclusion or removal of plant height at 450, 500, 550, and 600 GDD did not perform significantly different from the chosen model. For growth rate, 550 to 700 and 950 GDD had the lowest partial R^2^ values as well as some of the lowest relative importance during model selection with growth rate of 450, 600, and 950 having lower weights and 600 GDD removed from the model completely. The chosen model for growth rate was the only model with a delta AIC lower than 2, which indicated it was significantly better than other models, though the removed time points did not consistently align with the time points of the weakest correlations with yield (Figure 3A). This analysis overall demonstrated limited explanatory information from very early plant height and growth rate for end of season grain yield.

### End of season grain yield can be predicted from vegetative growth curves

Due to the likelihood for many small and non-linear relationships between predictors and outcomes, as well as to have a generalized model, machine learning models were employed to predict grain yield from mid-season traits. Six models (lasso, pcaNNet, pls, plsRglm, rf, svmRadial, and svmPoly) were initially trained and tested with all data features (Table 1; Supplemental Table S7). After initial testing, three models (pls, plsRglm, and rf) were used for further testing based on the initial model performances. The three modeling techniques were trained using various morphological and developmental feature combinations that included temporal plant height, temporal growth rate, information about the exponential growth period, and FPCA scores derived from the full growth curves (i.e. growth eigen) (Table 1). The data was partitioned into training and test sets by year to prevent data leakage between the two sets, which resulted in three training and test sets (training with 2020 and 2021 and testing with 2022, training with 2020 and 2022 and testing with 2021, and training with 2021 and 2022 and testing with 2020). The highest performing hyperparameter values for each machine learning model-by-data type combination were compared, and the plsRglm model performed the highest, with an average Pearson correlation of 0.398 and an average root mean square error (RMSE) of 39.0. Plant height and growth rate (r = 0.344-0.423) features together showed greater success when used to predict grain yield than growth eigen alone (r = 0.198-0.393), and the addition of growth eigen to plant height and growth rate did not always improve model performance (Table 2). When plant height, growth rate, and growth eigens were all included in a single plsRglm model, the best average r (0.426) and RMSE (38.5) was observed with fairly high Pearson correlations in 2020 and 2022 (r = 0.697 & 0.568).

**Table 2.**
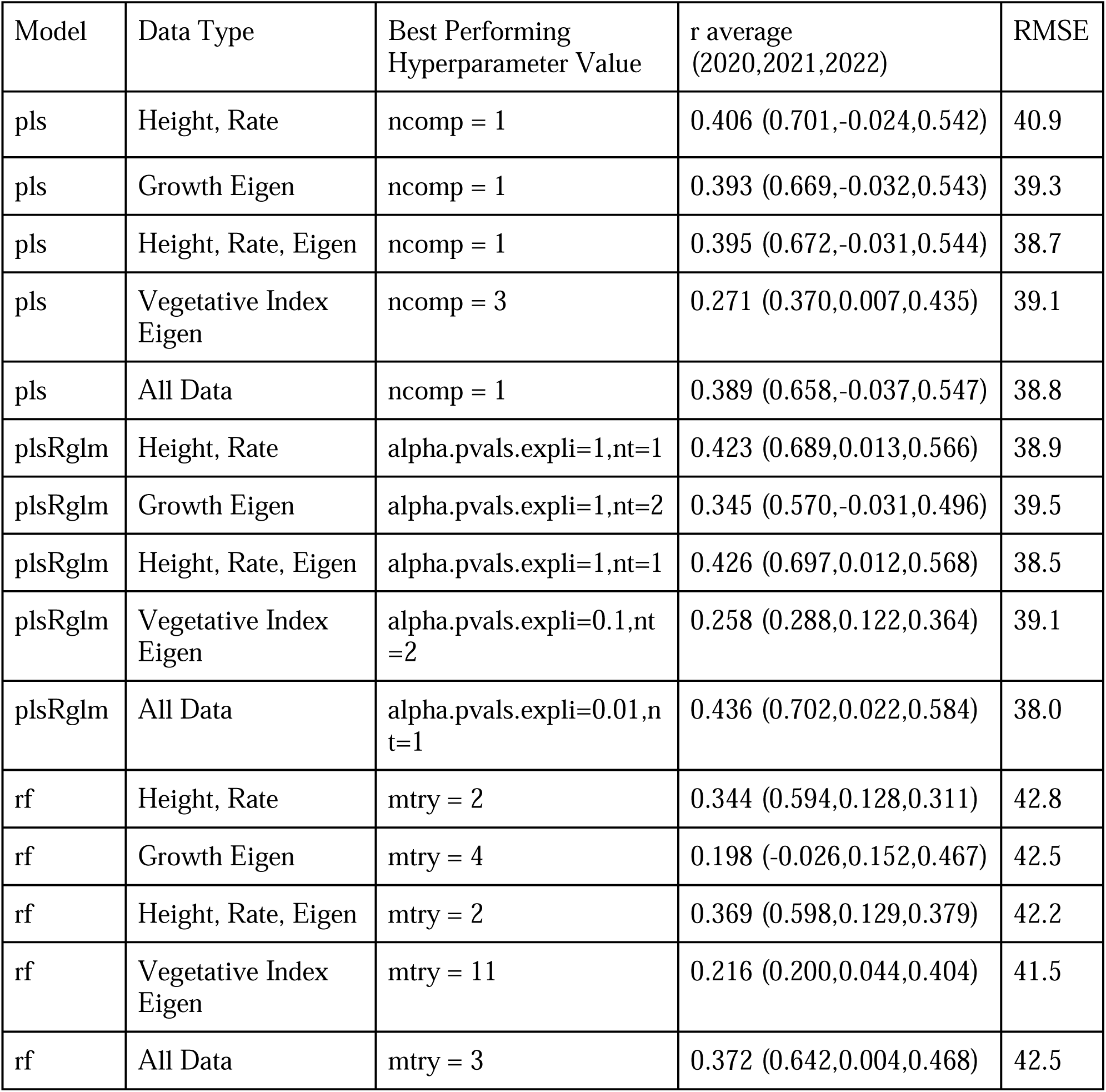
Average performance of model and data type combinations. Each model and data type is associated with the best performing hyperparameter value, the average r value, and the average RMSE across each testing and training set combination. Individual r values are also included for separate iterations with the year representing the specific year left out during model training and used as the test set.

In addition to morphological features, vegetative indices were also extracted from the UAV collected RGB images (Supplemental Table S6). Vegetative indices are linked to different plant physiological features that could be beneficial for predicting end of season grain yield. For instance, the normalized green red difference index has shown a strong relationship with chlorophyll and biomass (Tucker, 1979), both of which have been correlated with grain yield (Liu et al., 2019; Li et al., 2023). RGB derived vegetation indices have been successful at predicting maize yield on research plots (Adak et al., 2021), but have yet to be tested in commercial fields. A challenge in using vegetative indices in modeling across years is that they can not be modeled as a continuous curve from which predictors can be extracted at regular intervals throughout the growing season. Instead, to get comparable values across years with different flight dates, FPCA was performed across the flight dates within each year (Adak et al., 2021). Yield prediction using vegetation indices alone did not perform as well as was expected with r values ranging from 0.216 to 0.271. However, adding the vegetative indices to all the other data features did modestly improve overall Pearson correlation values from an average of 0.397 with only morphological and developmental data to 0.399 with vegetative indices included. Although the improvement in model performance was less than previously observed (Adak et al., 2023). This is likely due to the highly variable flight collection time points between years, which was not a factor in previous studies applying this approach in only a single growth environment (Adak et al., 2023). While a FPCA was used to make these data points more comparable over years, the lack of consistency and number of time points in a non-continuous trait made the comparison difficult. The accuracy of the vegetative indices collected may have also been impacted by the lack of atmospheric calibration. Due to the size of the fields, vegetative indices collected in the same flight may even have variation in effects due to atmospheric changes over the course of the flight. The inclusion of vegetative indices could be more helpful if the flights were conducted at more comparable timepoints and with calibration targets included in the images. Still, the plsRglm model trained with all data features, which included vegetative indices, performed the best across all three years (Figure 4A and 4B) with an average Pearson correlation of 0.436 and the lowest average RMSE values at 38.0 (Table 2).

**Figure 4.**
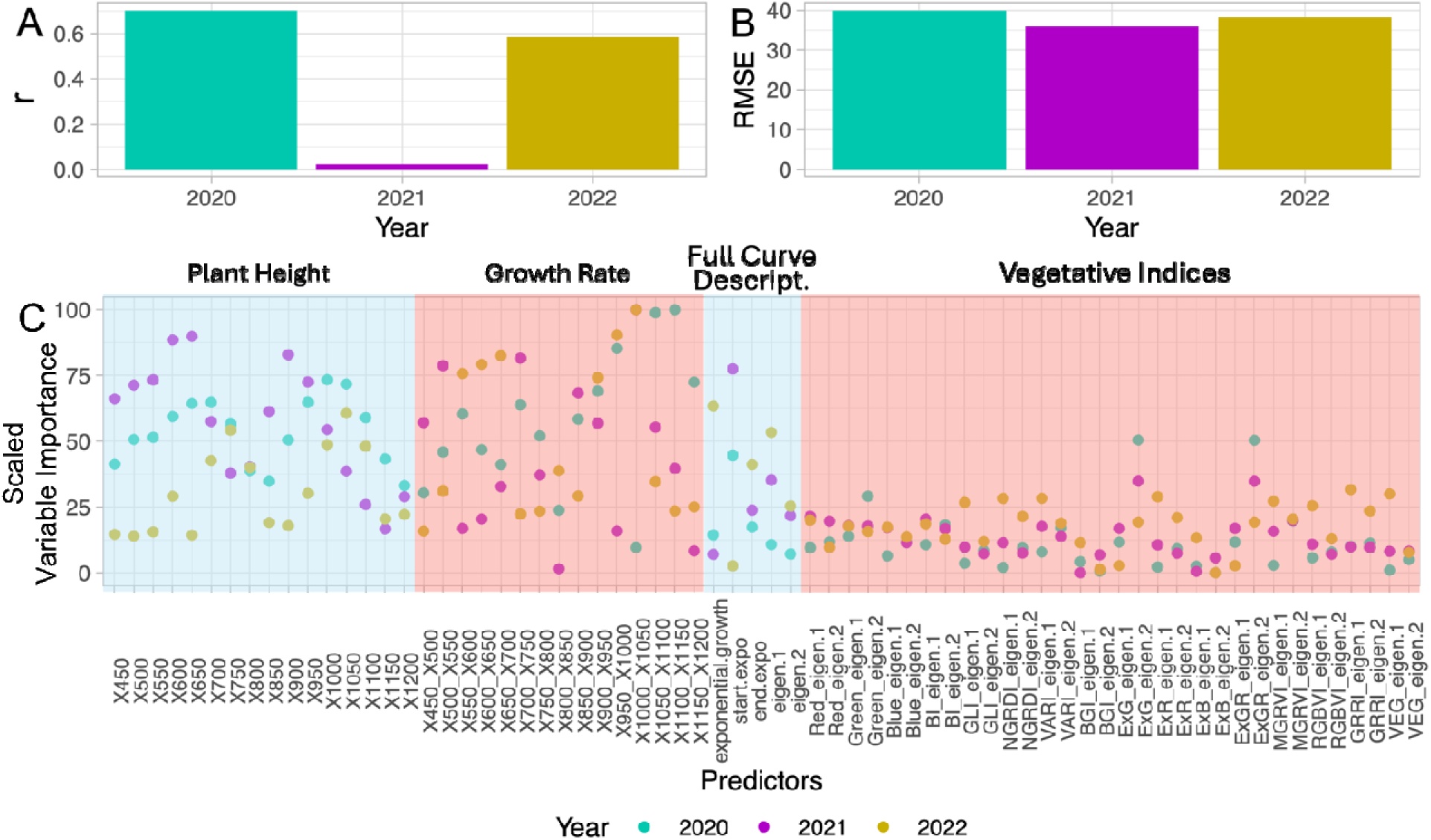
Prediction of grain yield with a plsRglm model trained using plant height, growth rate, and vegetative index features. (A) Pearson correlation coefficient (r) of predicted average yield compared to actual average yield within plots across different sets of training and test data (e.g. 2020 indicates predictions on 2020 data based on a model trained with 2021 and 2022 data). (B) Root Mean Square Error (RMSE) of predicted average yield when compared to actual average yield within plots. (C). Scaled variable importance of features included in the final machine learning model.

To determine the relative importance of the different features included in the plsRglm models the scaled variable importance of each combination of training years was evaluated. No data information was used exactly the same across all three combinations of years, but some trends were observed (Figure 4C). Overall, plant height and growth rate had higher scaled variable importance compared to any of the FPCA scores associated with the full growth curve or the vegetative indices. This makes sense given that the Pearson correlations from using the plant height and growth rate data for model training was higher (r = 0.344-0.423) than when the model was trained using the full growth curve eigenvalues or the vegetative indice FPCA scores (r = 0.198-0.393; Table 2). Still, some vegetative indices contributed to the overall performanc of the model, with ExG and ExGR both showing relatively high importance in predicting grain yield for 2020 and 2021 compared to the other vegetative indices (Figure 4C). ExG was originally developed to separate crop materials from weeds, background noise, and dead plant material (D. M. Woebbecke et al., 1995). Other studies have confirmed the usefulness of ExG to identify plant material as well as functionality in predicting the Normalized Difference Vegetation Index (Larrinaga and Brotons, 2019; Mehrotra and Srinivasan, 2019), which can be used to monitor crop health and productivity using near-infrared wavelengths (Johnson et al., 2021). ExGR has similarly been shown to identify healthy plant material and has a strong relationship with ExG (Mehrotra and Srinivasan, 2019; Lu et al., 2019).

### Water sufficiency influenced the ability of growth curves to predict yield

Predicting yield using machine learning resulted in fairly similar performance for 2020 and 2022, but the 2021 predictions had a much lower correlation to measured yields no matter the data combination or modeling approach used (Supplemental Figure S6, Supplemental Table S7). The low variation in plant height and growth rates within each time point in 2021 may be a factor in the low modeling accuracy (Figure 1D). Furthermore, the shape of the growth curves in 2021 was also quite different from 2020 and 2022 (Figure 1D), which may have also contributed to the poor model performance when predicting 2021 data as differently shaped growth curves would be outside the training boundaries of the model.

We wanted to further understand why there was a large difference in growth curves in 2021, and hypothesized some of the variation could be due to differences in weather. The precipitation in 2021 was much lower than in either 2020 or 2022 (10.89 inches in 2020, 6.41 inches in 2021, 10.30 inches in 2022 from planting to flowering), and 2021 accumulated substantially more solar radiation (Supplemental Figure S7). In a drought year, such as was the case in 2021, yield can be greatly affected by soil type (Butts-Wilmsmeyer et al., 2019), and specifically its ability to hold water, as well as by the depth to the water table (Kadioglu et al., 2019; Odili et al., 2023). To test if soil type and water table depth were contributing factors in this study, we acquired soil maps from the National Resource Conservation Service website to identify rough outlines of soil type and depth to water table differences across the fields (National Cooperative Soil Survey, 2019). Overlaying the soil map on the yield maps showed little relationship with yield in 2020 or 2022 (Figure 5A and C), but indicated a possible connection between soil type and yield in 2021 (Figure 5B). The lowest yielding area in 2021 was in the bottom center and bottom left areas of the field (Figure 5B - I), which contained Nicollet clay loam with 1 to 3 percent slopes. This area was identified as having a 12 to 24 inch depth to the water table with somewhat poorly drained soil. The top left area of the field (Figure 5B - V) also had lower yields where the soil was moderately eroded Clarion loam with 6 to 10 percent slopes. This soil was defined as well drained with a depth to water table of 47 to 63 inches. Unlike both of these areas, the highest yielding portion of the field (Figure 5B - VI) contained Delft, overwash-Delft complex soil with 1 to 4 percent slopes. The depth to the water table in this area of the field is only 6-18 inches and the soil is poorly drained to somewhat poorly drained, and therefore not only has a shallower depth to the water table, but is able to retain precipitation better. The same field that was flown in 2021 was also flown in 2022. In 2022, there was little to no connection between yield and soil type/depth maps, which indicated that yield is not as influenced by soil type or depth to the water table when there is sufficient precipitation throughout the growing season.

**Figure 5.**
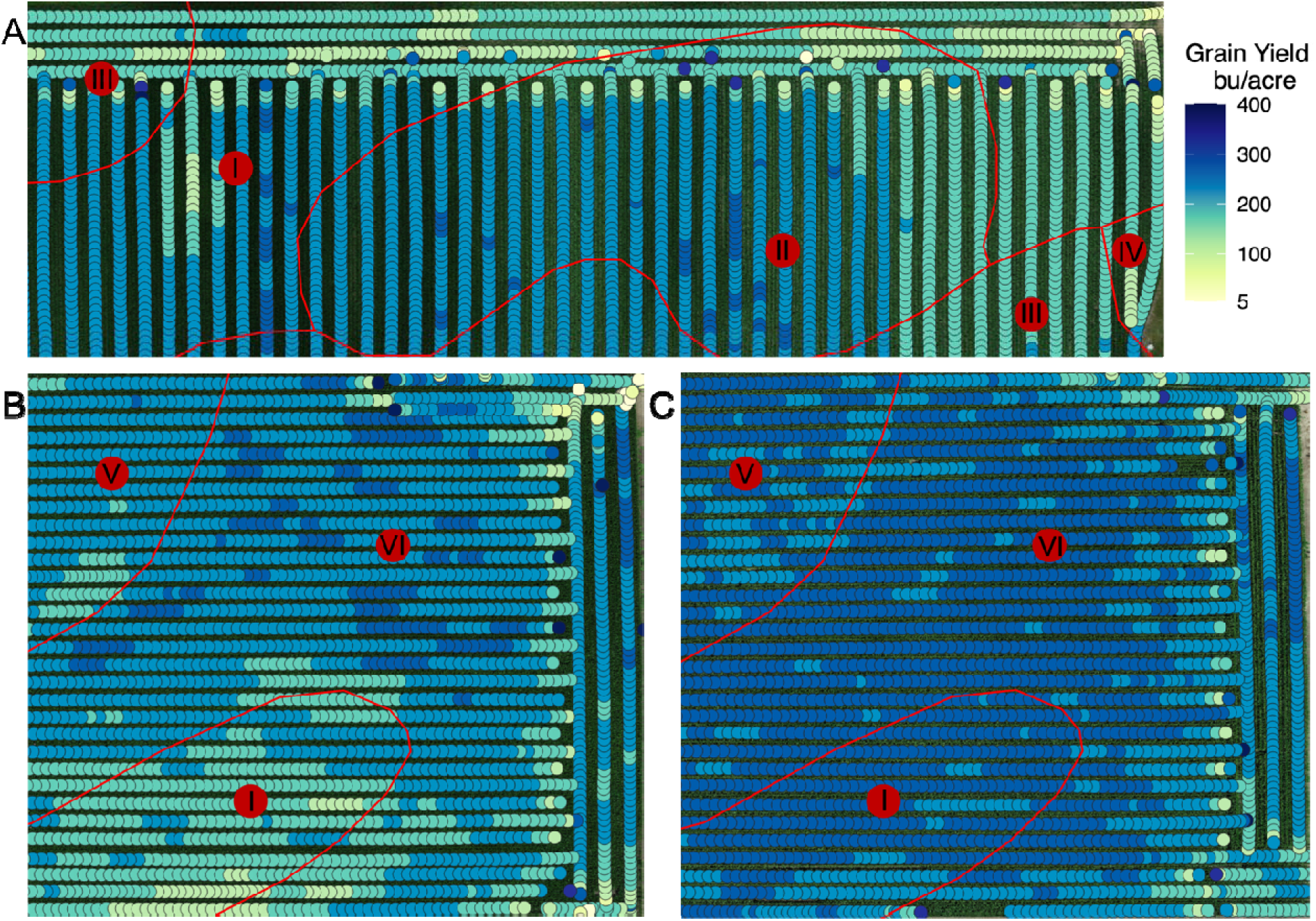
Overlay of soil maps on grain yield maps. Grain yield maps (circle data points) with soil map location layovers (red outlined areas with numbers denoting different soil types within a field) for (A) 2020, (B) 2021, and (C) 2022. (I) Nicollet clay loam, 1 to 3 percent slopes, (II) Clarion loam, 2 to 6 percent slopes, (III) Webster clay loam, 0 to 2 percent slopes, (IV) Canisteo-Glencoe complex, 0 to 2 percent slopes, (V) Clarion loam, 6 to 10 percent slopes, moderately eroded, (VI) Delft-overwash Delft complex, 1 to 4 percent slopes.

These results collectively indicate that, in cases of sufficient water, growth curves can be effective indicators of end of season yield and have great potential to contribute to management strategies used in precision farming. However, without sufficient water, growth curves may not provide as useful information for management strategies, and dependence on soil type and depth to water tables may be more useful under these conditions.

## Conclusion

The results of this study have shown the potential for early season within-field yield estimation in production fields using RGB image extracted values. Further data collection from more fields across more years will improve this method of yield estimation and could potentially make it possible to optimize models for soil type, depth to water table, and weather conditions of each field in each growing year. For instance, predicting yield in 2021 showed little success, but there were no other drought years to train the model. Thus, while there were clear relationships between vegetative plant height, growth rate, and vegetative indices with end of season yield, to realize the full potential of this approach information on substantially more growth environments will be required.

## Acknowledgements

We would like to thank Amanda Gilbert for technical support for this experiment. We would also like to thank the Minnesota Supercomputing Institute at the University of Minnesota (http://www.msi.umn.edu) for providing resources that contributed to the research results reported in this article. This work was supported by the Minnesota Corn Research and and Promotion Council. DDS was funded by the University of Minnesota MnDRIVE Global Food Ventures Graduate Fellowship. JC was funded by the Bayer/University of Minnesota Multifunctional Agriculture Initiative Graduate Student Fellowship.

## Abbreviations

AIC: Akaike Information Criterion
DSM: Digital Surface Model
FPCA: Functional Principal Component Analysis
GCP: Ground Control Point
GDD: Growing Degree Day Units
RGB: Red Green Blue
RMSE: root mean square error
UAV: Unoccupied Aerial Vehicle

