## Supplemental Figures 1-7 for "Plant height defined growth curves during vegetative development have the potential to predict end of season maize yield and assist with mid-season management decisions"

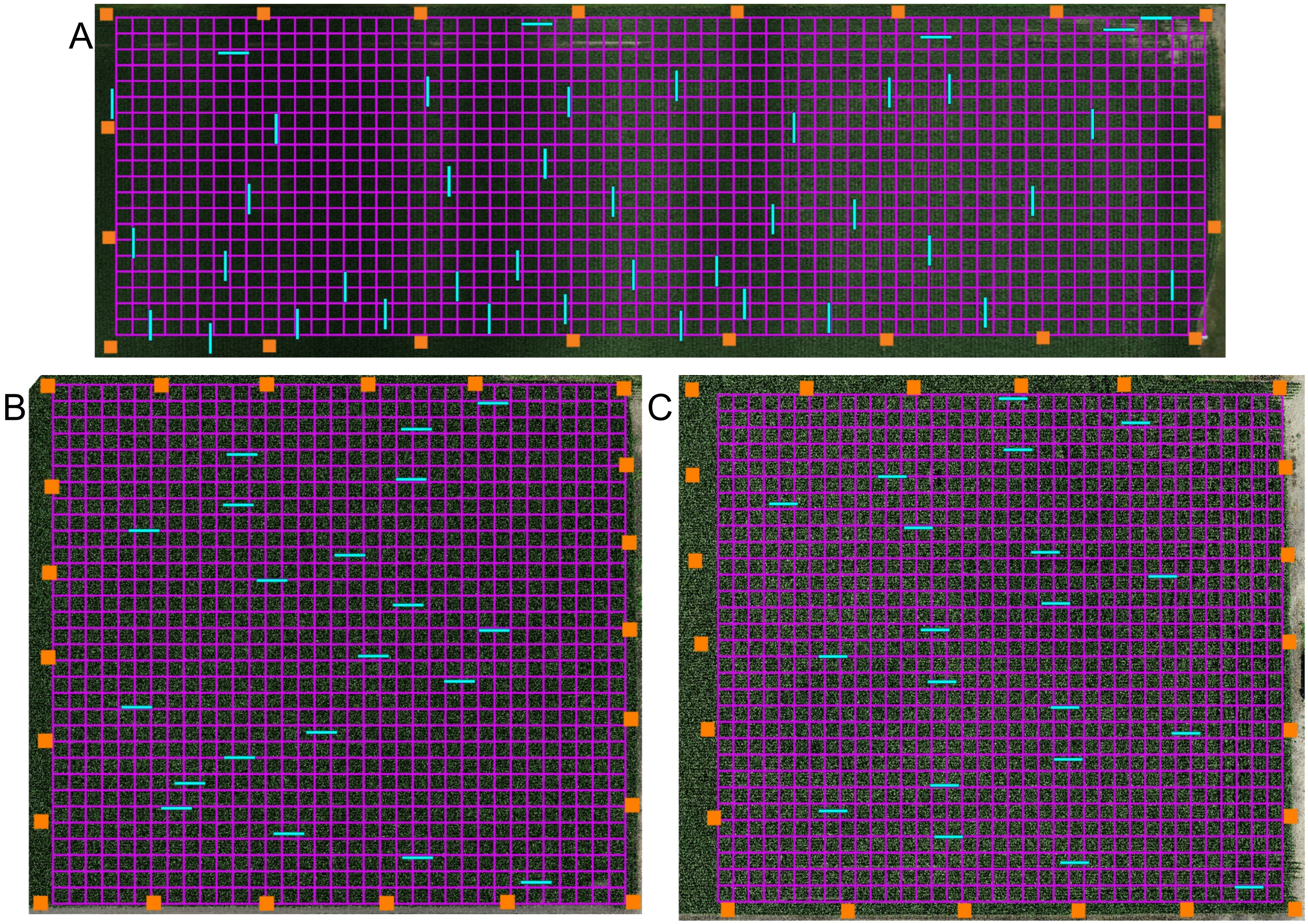


**Supplemental Figure S1.** Plot extraction boundaries and location of ground control points. The plot boundaries for plant height extraction in (A) 2020, (B) 2021, and (C) 2022. The pink grid shows the plot boundaries across the whole field, the blue plot boundaries were the manually measured plots, and the orange squares show the location of the ground control points.

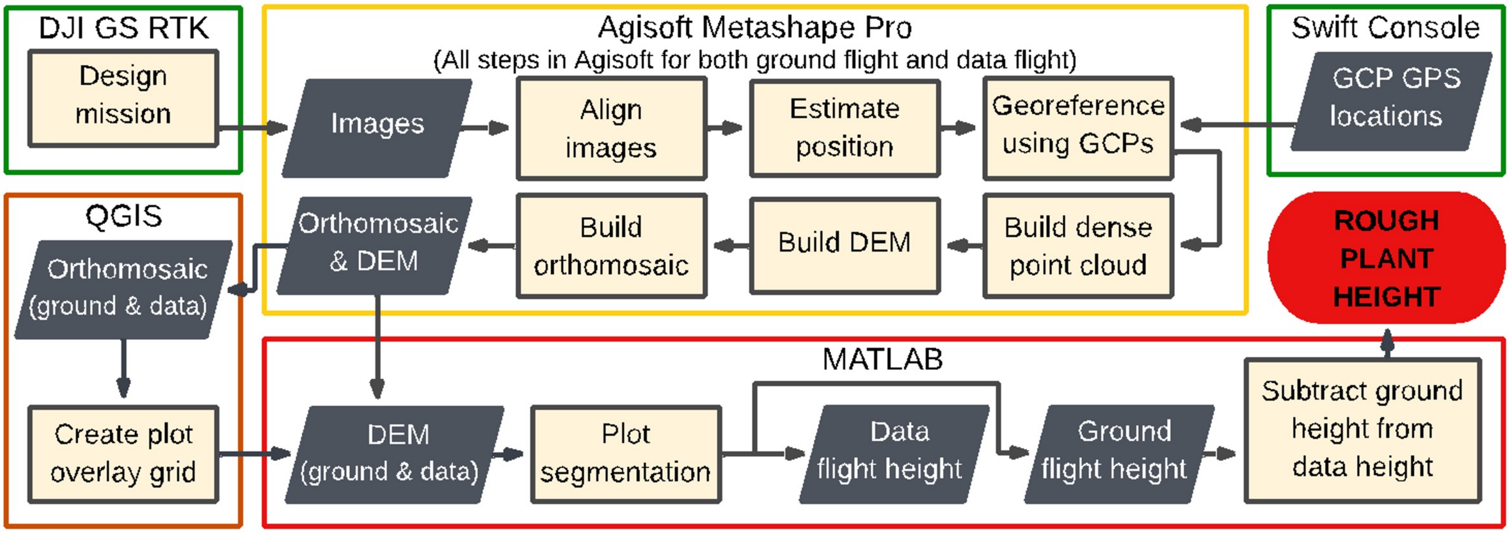


**Supplemental Figure S2.** Unoccupied aerial vehicle data processing and plant height extraction pipeline.

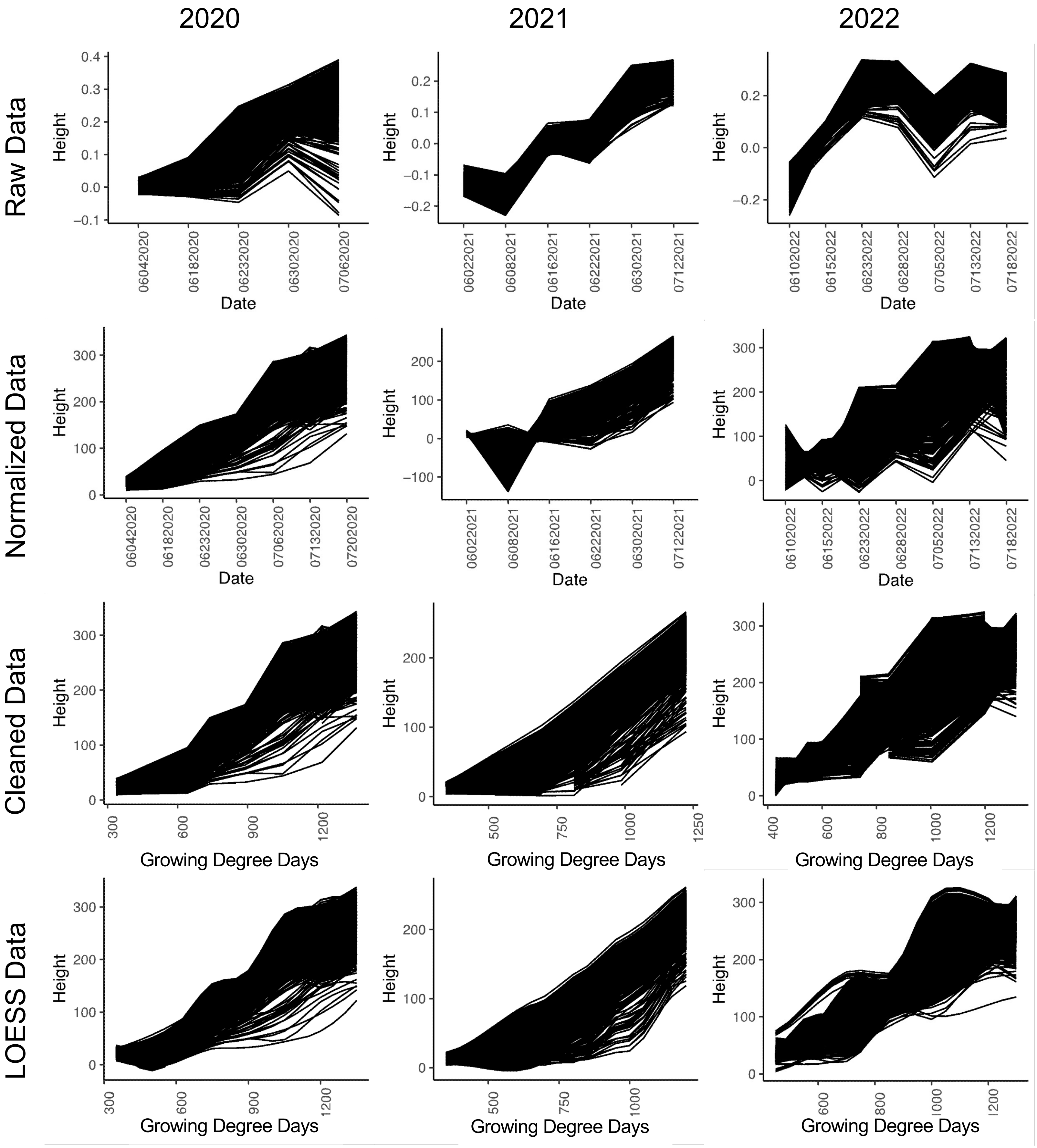


**Supplemental Figure S3.** Normalization of extracted plant height values. Extracted plant height values for all plots with raw data before any cleaning (Raw Data), before cleaning but normalized across dates using ground control point heights (Normalized Data), normalized data with erroneous plots removed before any analysis (Cleaned Data), and LOESS curves fit to the extracted cleaned extracted data (LOESS Data). Dates on the x-axis are in the form MMDDYYYY for Raw and Normalized Data and in Growing Degree Days (˚F) for Cleaned and LOESS Data.

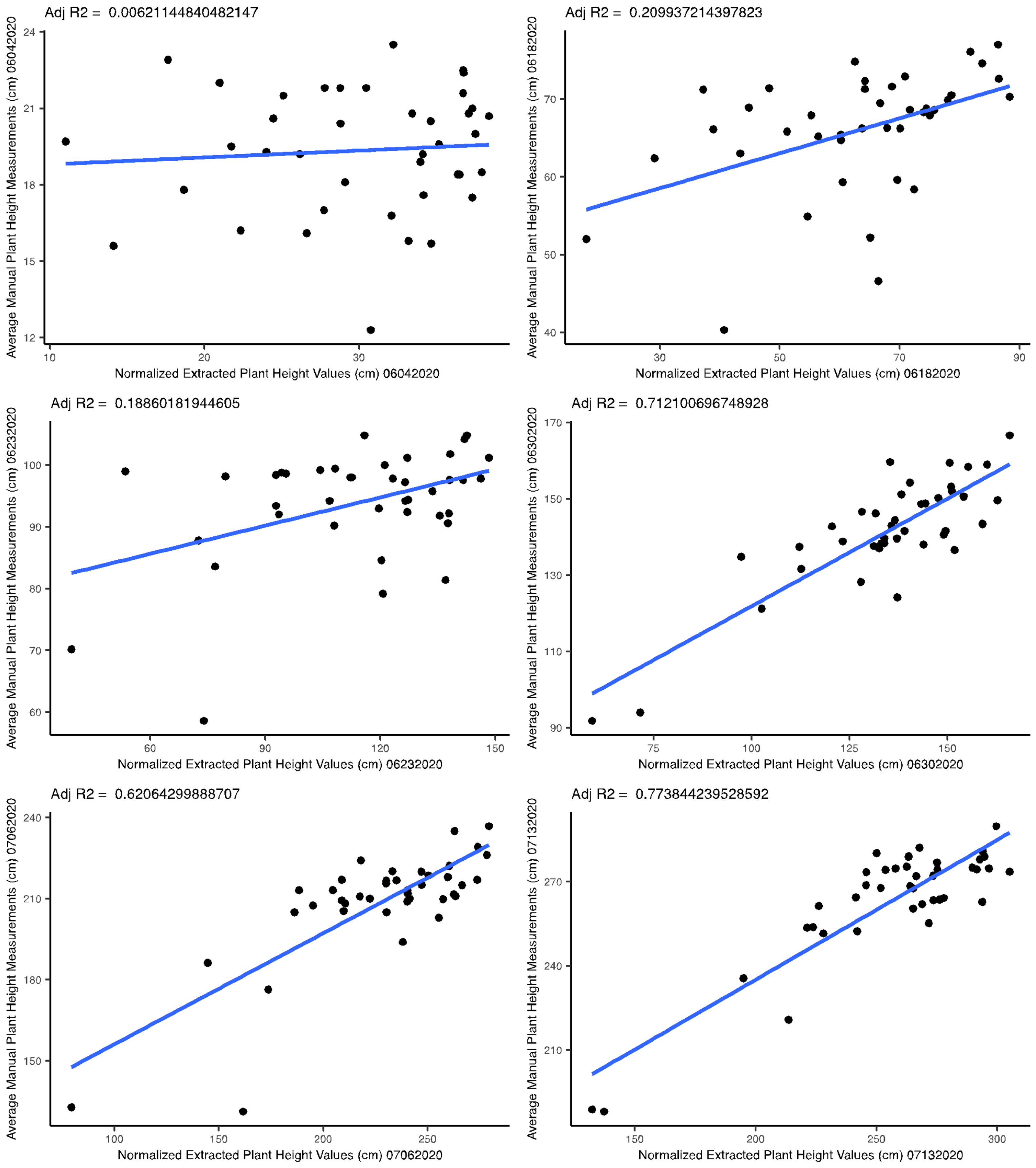

Adj R2 = 0.00621144840482147
Average Manual Plant Height Measurements (cm) 06042020
Normalized Extracted Plant Height Values (cm) 06042020
Adj R2 = 0.209937214397823
Average Manual Plant Height Measurements (cm) 06182020
Normalized Extracted Plant Height Values (cm) 06182020
Adj R2 = 0.18860181944605
Average Manual Plant Height Measurements (cm) 06232020
Normalized Extracted Plant Height Values (cm) 06232020
Adj R2 = 0.712100696748928
Average Manual Plant Height Measurements (cm) 06302020
Normalized Extracted Plant Height Values (cm) 06302020
Adj R2 = 0.62064299888707
Average Manual Plant Height Measurements (cm) 07062020
Normalized Extracted Plant Height Values (cm) 07062020
Adj R2 = 0.773844239528592
Average Manual Plant Height Measurements (cm) 07132020
Normalized Extracted Plant Height Values (cm) 07132020

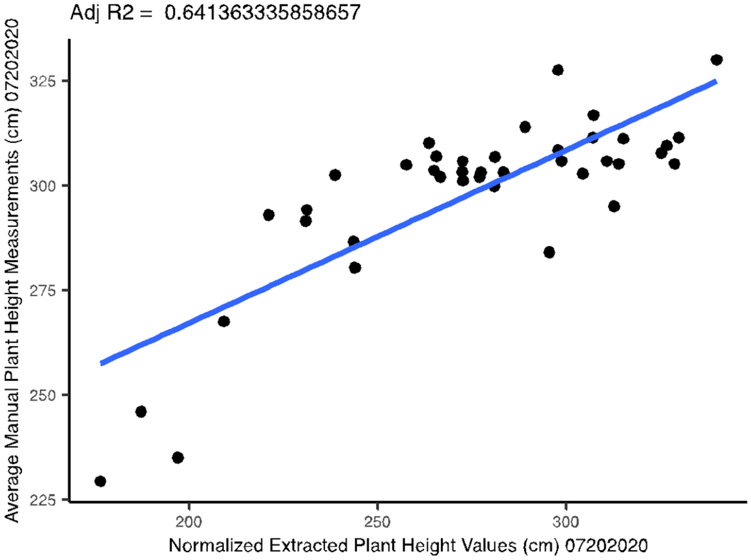
Adj R2 = 0.641363335858657
Average Manual Plant Height Measurements (cm) 07202020
225
250
275
300
325
200
250
300
Normalized Extracted Plant Height Values (cm) 07202020

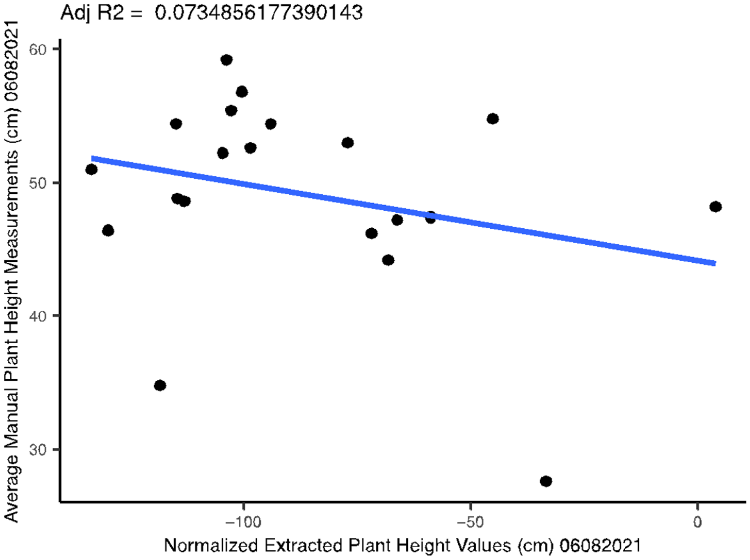
Adj R2 = 0.0734856177390143
Average Manual Plant Height Measurements (cm) 06082021
30
40
50
60
−100
−50
0
Normalized Extracted Plant Height Values (cm) 06082021

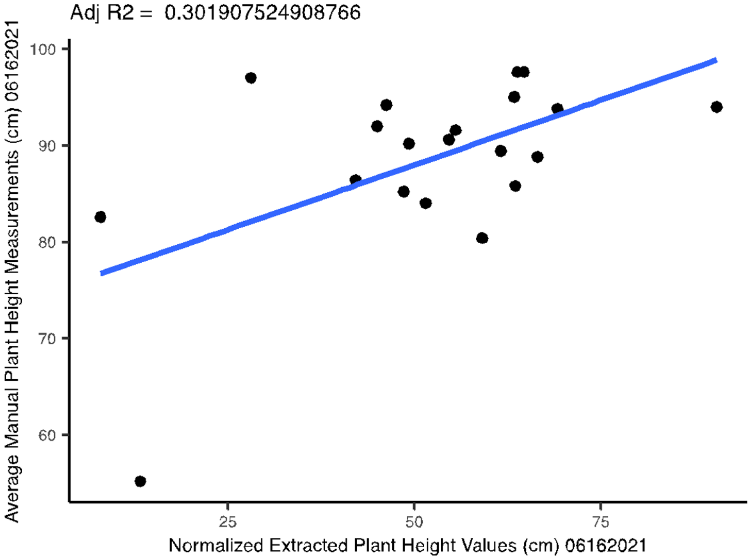
Adj R2 = 0.301907524908766
Average Manual Plant Height Measurements (cm) 06162021
60
70
80
90
100
25
50
75
Normalized Extracted Plant Height Values (cm) 06162021

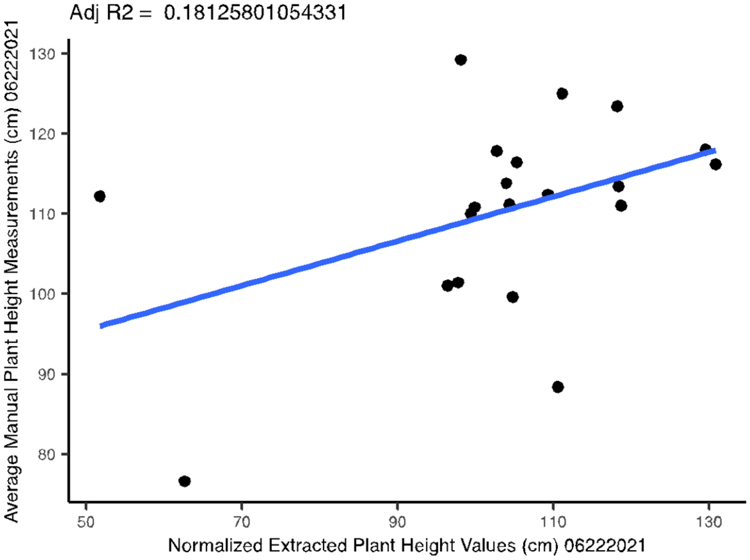
Adj R2 = 0.18125801054331
Average Manual Plant Height Measurements (cm) 06222021
80
90
100
110
120
130
50
70
90
110
130
Normalized Extracted Plant Height Values (cm) 06222021

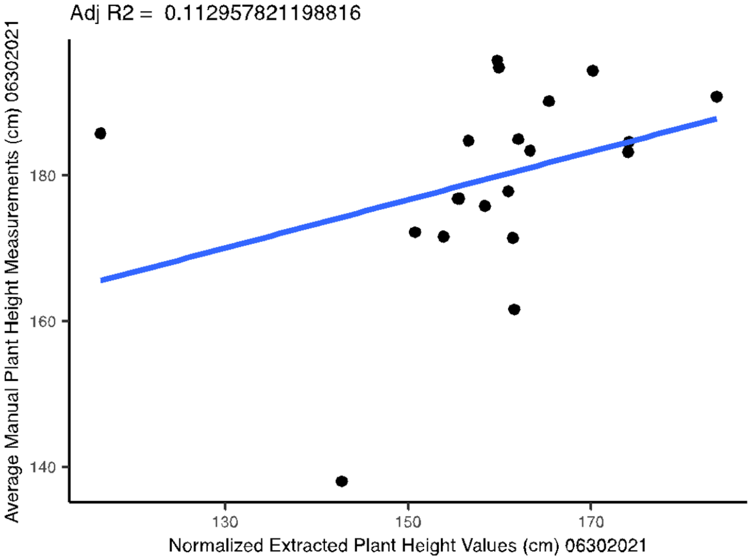
Adj R2 = 0.112957821198816
Average Manual Plant Height Measurements (cm) 06302021
140
160
180
130
150
170
Normalized Extracted Plant Height Values (cm) 06302021

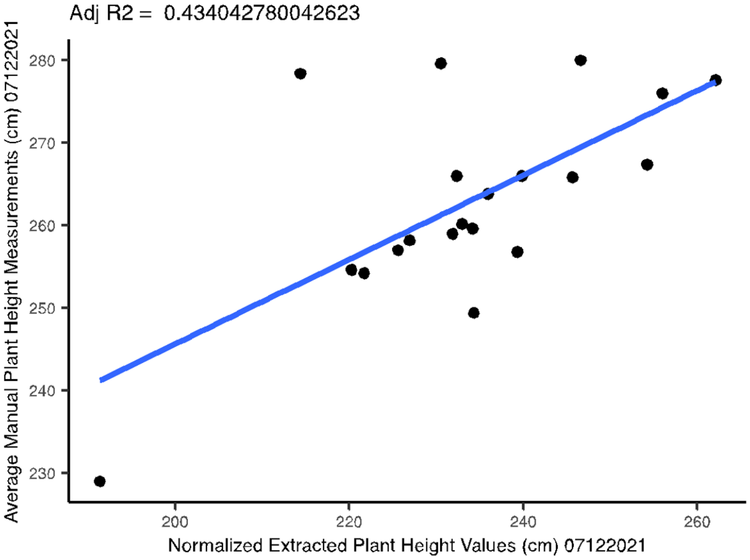
Adj R2 = 0.434042780042623
Average Manual Plant Height Measurements (cm) 07122021
230
240
250
260
270
280
200
220
240
260
Normalized Extracted Plant Height Values (cm) 07122021

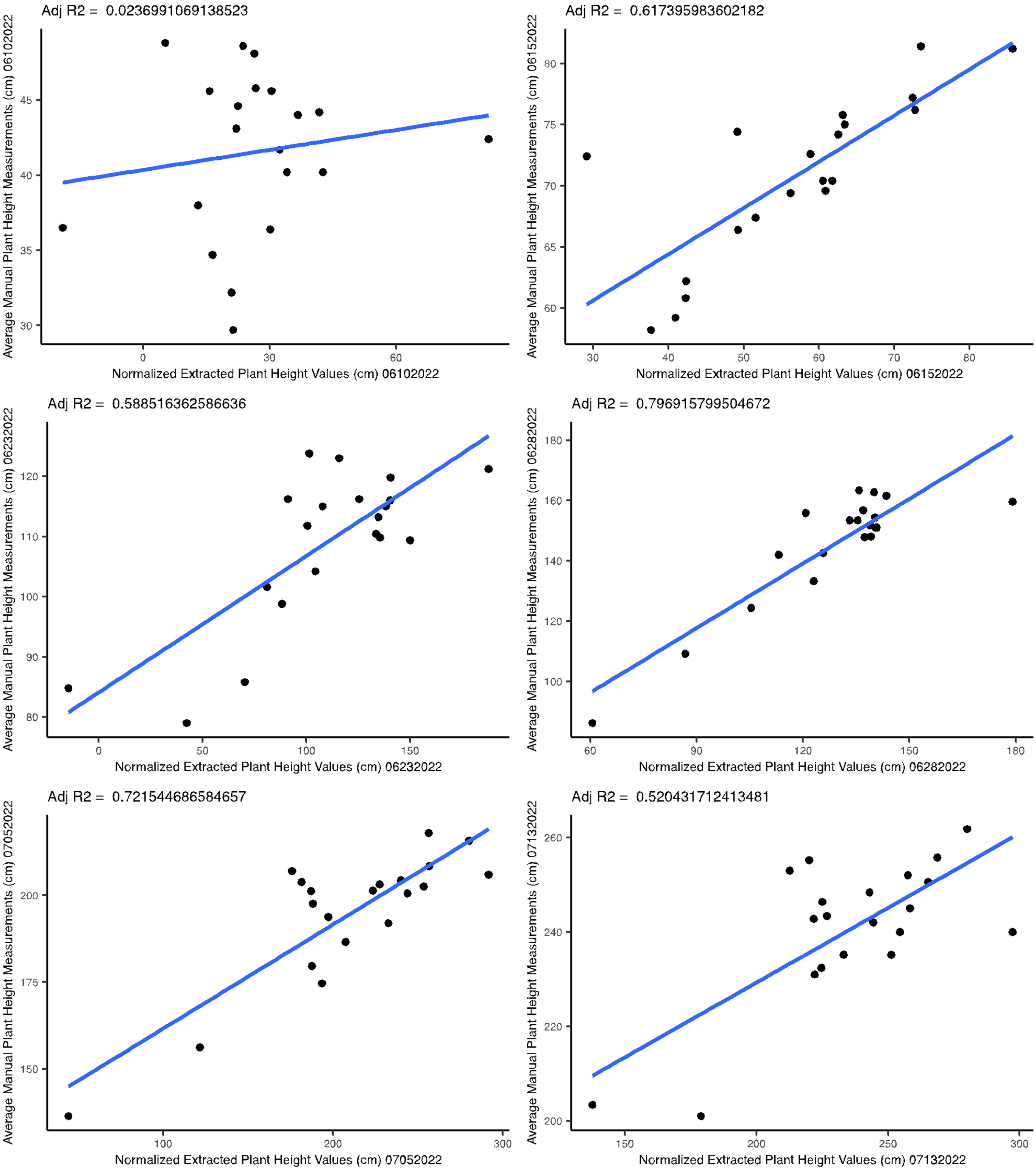
Adj R2 = 0.0236991069138523
Average Manual Plant Height Measurements (cm) 06102022
Normalized Extracted Plant Height Values (cm) 06102022
Adj R2 = 0.617395983602182
Average Manual Plant Height Measurements (cm) 06152022
Normalized Extracted Plant Height Values (cm) 06152022
Adj R2 = 0.588516362586636
Average Manual Plant Height Measurements (cm) 06232022
Normalized Extracted Plant Height Values (cm) 06232022
Adj R2 = 0.796915799504672
Average Manual Plant Height Measurements (cm) 06282022
Normalized Extracted Plant Height Values (cm) 06282022
Adj R2 = 0.721544686584657
Average Manual Plant Height Measurements (cm) 07052022
Normalized Extracted Plant Height Values (cm) 07052022
Adj R2 = 0.520431712413481
Average Manual Plant Height Measurements (cm) 07132022
Normalized Extracted Plant Height Values (cm) 07132022

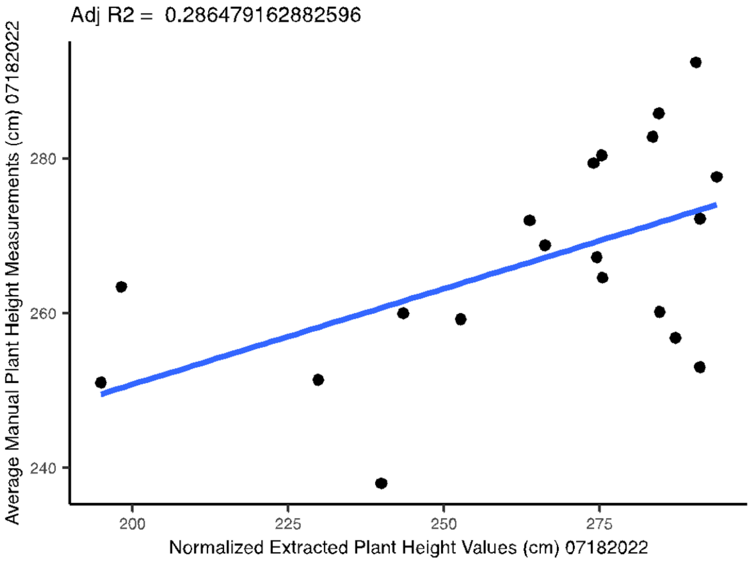


**Supplemental Figure S4.** Validation of extracted plant height. Pearson correlation of normalized mean extracted plot plant height to mean manual plot plant height measurements. The date of each flight is indicated in the x- and y-axis label in the form MMDDYYYY.

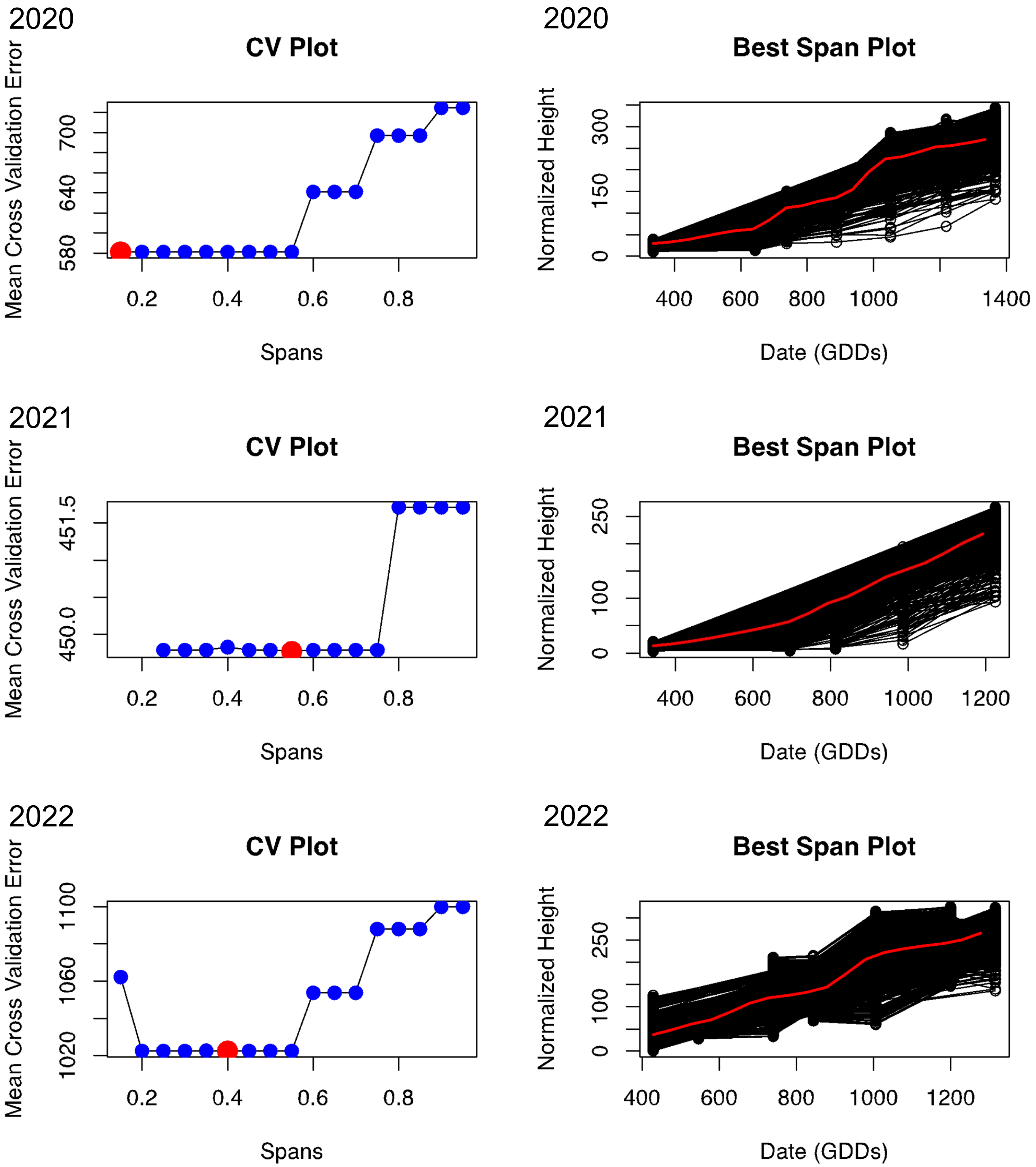


**Supplemental Figure S5.** Span choices for LOESS curve fitting each year. Mean cross validation error for each possible span from 0.15 to 0.95 for each year with the best span for each year shown with red dots in the left column plots. The growth rates with LOESS curves fitted using the best span are shown in the plots in the right column with the average curve across all plots in red.

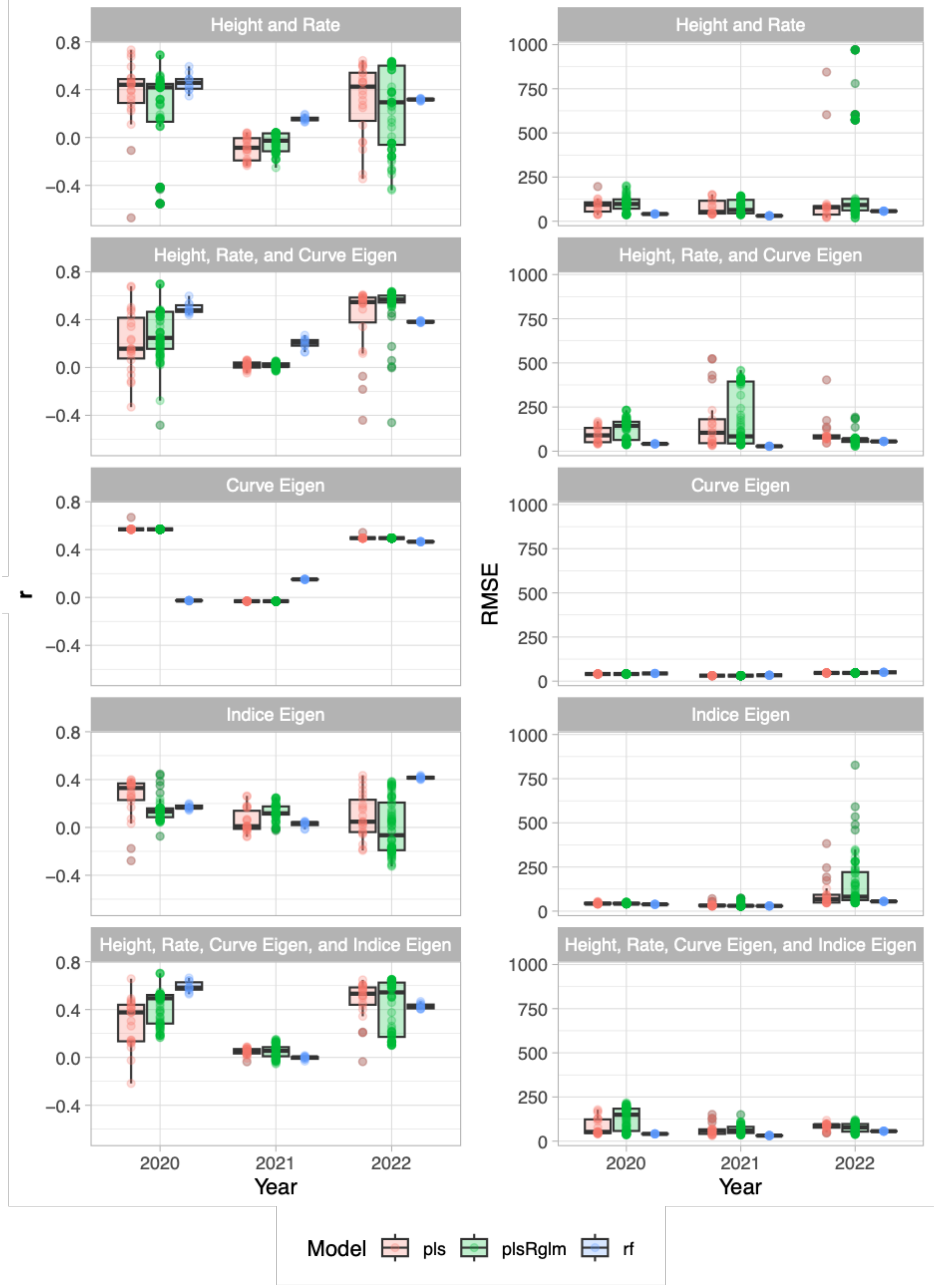


**Supplemental Figure S6.** Machine learning model performance. Pearson correlation coefficient (r) of predicted average yield compared to actual average yield within plots for each data combination (e.g. 2020 indicates predictions on 2020 data based on a model trained with 2021 and 2022 data), model, and data type (left). The plots in the right column are the root mean square error of predicted average yields when compared to actual average yield within plots for each data combination, model, and data type.

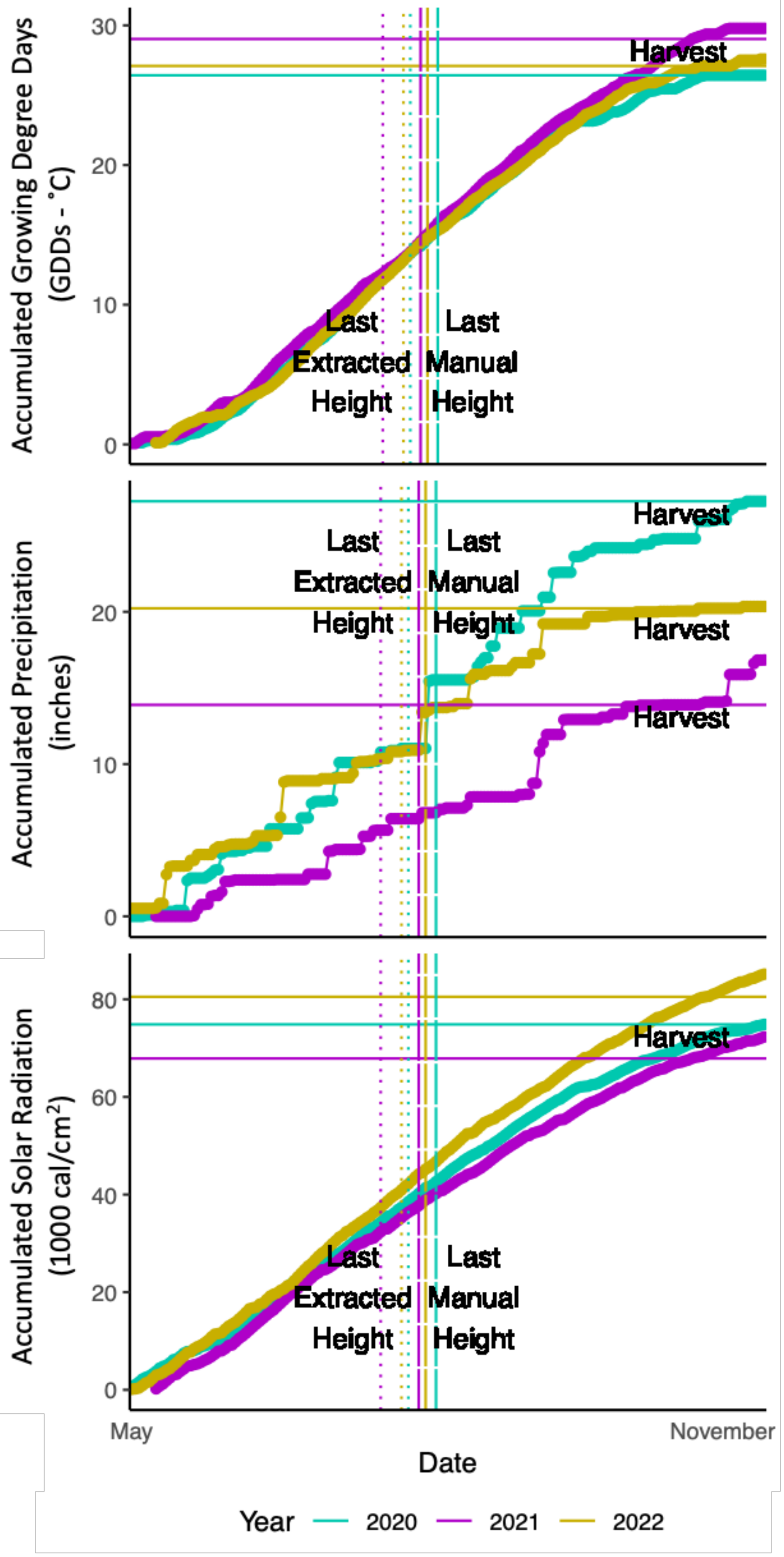


**Supplemental Figure S7.** Weather data for growing degree days, accumulated precipitation, and solar radiation accumulation across the growing seasons.
